# Decoding age-related changes in the spatiotemporal neural processing of speech using machine learning

**DOI:** 10.1101/786566

**Authors:** Md Sultan Mahmud, Faruk Ahmed, Rakib Al-Fahad, Kazi Ashraf Moinuddin, Mohammed Yeasin, Claude Alain, Gavin M. Bidelman

**Author notes:** **Address for editorial correspondence:** Md Sultan Mahmud, Department of Electrical and Computer Engineering, University of Memphis, 3815 Central Avenue, Memphis, TN, 38152.

## Abstract

Speech comprehension in noisy environments depends on complex interactions between sensory and cognitive systems. In older adults, such interactions may be affected, especially in those individuals who have more severe age-related hearing loss. Using a data-driven approach, we assessed the temporal (*when* in time) and spatial (*where* in the brain) characteristics of the cortex’s speech-evoked response that distinguish older adults with or without mild hearing loss. We used source montage to model scalp-recorded during a phoneme discrimination task conducted under clear and noise-degraded conditions. We applied machine learning analyses (stability selection and control) to choose features of the speech-evoked response that are consistent over a range of model parameters and support vector machine (SVM) classification to investigate the *time course* and *brain regions* that segregate groups and speech clarity. Whole-brain data analysis revealed a classification accuracy of 82.03% [area under the curve (AUC)=81.18%; F1-score 82.00%], distinguishing groups within ∼50 ms after speech onset (i.e., as early as the P1 wave).We observed lower accuracy of 78.39% [AUC=78.74%; F1-score=79.00%] and delayed classification performance when the speech token were embedded in noise, with group segregation at 60 ms. Separate analysis using left (LH) and right hemisphere (RH) regions showed that LH speech activity was better at distinguishing hearing groups than activity measured over the RH. Moreover, stability selection analysis identified 13 brain regions (among 1428 total spatiotemporal features from 68 regions) where source activity segregated groups with >80% accuracy (clear speech); whereas 15 regions were critical for noise-degraded speech to achieve a comparable level of group segregation (76% accuracy). Our results identify two core neural networks associated with complex speech perception in older adults and confirm a larger number of neural regions, particularly in RH and frontal lobe, are active when processing degraded speech information.

## 1. INTRODUCTION

Hearing impairment (HI) is the fifth leading disability worldwide (Vos et al., 2015) and the third most common chronic disease behind heart disease and arthritis (Blackwell, Lucas, & Clarke, 2014; Liberman, 2017). It is one of the core contributors to the growing disability problem in the United States (Murray et al., 2013). In older adults, HI has been associated with poor cognitive health, social isolation, and loneliness (Diaz, Johnson, Burke, Truong, & Madden, 2018; Lin et al., 2013). Robust speech processing in the elderly relies on a complex network of interacting brain regions (Bidelman, Mahmud, et al., 2019). Age-related HI is thought to occur due to a myriad of changes in both the central auditory pathways (Bidelman, Price, Shen, Arnott, & Alain, 2019; Bidelman, Villafuerte, Moreno, & Alain, 2014) and widespread areas of both cerebral hemispheres (Gates & Mills, 2005). For example, studies have shown aged-related declines in the temporal precision (Roque, Karawani, Gordon-Salant, & Anderson, 2019) of (subcortical) neural encoding (Anderson, Parbery-Clark, White-Schwoch, & Kraus, 2012; Bidelman et al., 2014; Konrad-Martin et al., 2012; Schoof & Rosen, 2016) and functional magnetic resonance imaging (fMRI) has shown older adults have greater activation than younger adults in widespread cortical brain regions (Diaz et al., 2018; Mudar & Husain, 2016). Older adults with hearing impairment show even greater activation in right hemisphere (RH) than the left hemisphere (LH) during speech perception in noise (Mudar & Husain, 2016).

Speech-in-noise (SIN) comprehension can be difficult for older adults, especially in those with hearing loss. The neurophysiological factors that influence SIN comprehension are not fully understood, but likely depends on rapid temporal processing. As such, tracking the neural encoding of speech processing necessitates use of neuroimaging techniques with excellent temporal resolution, such as event-related potentials (ERPs). EEG/ERPs also offer a non-invasive means for clinical diagnostics, including those related to cognitive aging as well as tracking how the brain encodes important features of the speech signal (Bidelman, Lowther, Tak, & Alain, 2017). For instance, the auditory cortical ERPs, comprised of the P1, N1, and P2 waves, are highly sensitive to the acoustic features of speech (Agung, Purdy, McMahon, & Newall, 2006), and correlate with listeners perception for both clear and degraded speech (Bidelman et al., 2014; Ross et al., 2009; Tremblay, Kraus, McGee, Ponton, & Otis, 2001).

Evidence suggests that older adults incorporate more attentional resources than younger adults in auditory perceptual tasks (Alain, McDonald, Ostroff, & Schneider, 2004; Grady, 2008). This could account for some of the age-related increases in ERP amplitudes reported in HI vs. NH listeners (Alain, 2014; Bidelman, Price, et al., 2019). Prior neuroimaging studies (Erb & Obleser, 2013; Peelle, Troiani, Wingfield, & Grossman, 2009; Vaden, Kuchinsky, Ahlstrom, Dubno, & Eckert, 2015; Wong et al., 2009) have also demonstrated increased activity in prefrontal regions related to cognitive control, attention, and working memory when older listeners process speech under challenging situations. Du and colleagues (Du, Buchsbaum, Grady, & Alain, 2016) demonstrated that older adults’ increased recruitment of frontal regions to perceive speech in noisy environments. In our earlier study (Bidelman, Mahmud, et al., 2019), we used functional connectivity analysis to demonstrate that older listeners with mild hearing loss had more extended (less integrated) communication pathways and less efficient information exchange across the brain than their normal-hearing peers; directed connectivity analysis further showed that age-related hearing impairment reverses the direction of neural signaling within important hubs of the auditory-linguistic-motor loop of the dorsal-ventral pathways (e.g., primary auditory cortex—inferior frontal gyrus— primary motor cortex), implying a rerouting of information within the same speech circuitry (Bidelman, Mahmud, et al., 2019). However, our previous study focused on a restricted set of speech-relevant brain regions compared to the widespread and distributed networks involved in speech-language function (Du, Buchsbaum, Grady, & Alain, 2014; Du et al., 2016; Rauschecker & Scott, 2009). Extending prior work, we used here a comprehensive machine learning approach to identify the most probable *global* set of brain regions that are sensitive to senescent changes in speech processing. To our knowledge, this is the first study to apply decoding and machine learning techniques to map spatiotemporal differences in speech processing in older listeners at the full-brain level.

The current study aimed to further investigate the neural correlates of hearing loss on full-brain functionality using a data driven multivariate approach (machine learning). In this study, we used a source montage to analyze scalp-recorded speech-evoked responses to (i) develop a robust model to characterize neural activities involved in speech perception and (ii) categorize hearing loss based on speech-ERPs in different acoustic environments (e.g., clear and noise conditions).

We hypothesized that speech-evoked responses among normal hearing (NH) and HI listeners differ with regards to time and spatial regions that are recruited during phoneme discrimination. We further hypothesized that the core speech network “decoded” via machine learning vary as a function of the clarity of the speech signal. We applied a data-driven approach (support vector machine (SVM), stability selection) to source-level EEG data to identify when (*in time*) and where *(brain regions of interest, ROIs*) hearing status could be decoded from speech-evoked responses. We used a sliding window decoder to address the temporal dynamics of decoding and identify *when* speech-evoked responses distinguished older adults with normal hearing of mild hearing loss. In addition, stability selection, a machine learning approach to identify highly consistent data features, was used to examine *where* in the brain group responses best separated older adults with normal or mild hearing impairment.

## 2. MATERIALS & METHODS

Analyses of the ERPs and behavioral data associated with this dataset have been reported elsewhere (Bidelman, Mahmud, et al., 2019; Bidelman, Price, et al., 2019). In this study, we present a new machine learning analysis to identify the most discriminating spatiotemporal features of full-brain neuroelectric activity that best segregates normal and hearing-impaired listeners in terms of their SIN processing.

### 2.1 Participants

The sample consisted of thirty-two older adults (13 NH and 19 HI; aged 52-72 years). Demographic details are provided in our previous reports (Bidelman, Mahmud, et al., 2019; Bidelman, Price, et al., 2019; Mahmud et al., 2018). Listeners were divided into two cohorts based on their average hearing thresholds being better (NH) or poorer (HI) than 25 dB HL across both ears (Figure 1). The groups were matched in age (NH: 66.2±6.1 years, HI: 70.4±4.9 years; *t*_22.2_=−2.05, *p* = 0.052) and gender balance (NH: 5/8 male/female; HI: 11/8; Fisher’s exact test, *p*=0.47). Age and hearing loss were not correlated (Pearson’s *r*=0.29, *p*=0.10). All originally gave written informed consent in accordance with a protocol approved by the Baycrest Research Ethics Review Board.

**Figure 1:**
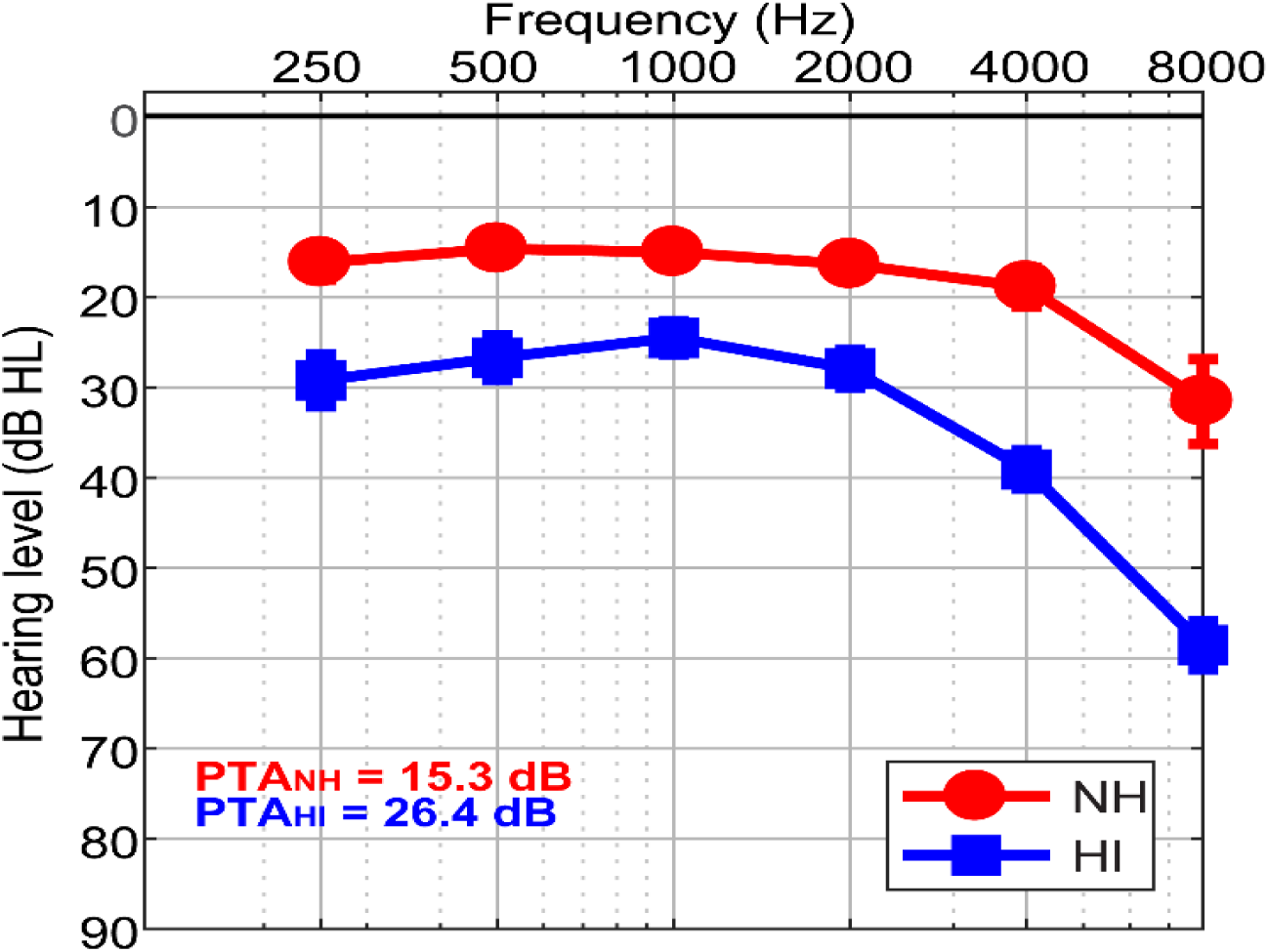
Behavioral audiograms (hearing thresholds) per group. NH, normal-hearing listeners, HI, hearing-impaired listeners.

### 2.2 Stimuli & task

Speech tokens (/ba/, /pa/, and /ta/; 100 ms) (Dubno & Schaefer, 1992) were presented back-to-back in random order with a jittered interstimulus interval (95-155 ms). Frequent (/ba/, /pa/; 3000 each) and infrequent (/ta/; 210) tokens were presented according to a pseudo-random schedule where at least two frequent stimuli intervened between target /ta/ tokens. Listeners were asked to detect target /ta/ tokens. For noise blocks, the speech triplet was mixed with noise babble (Killion, Niquette, Gudmundsen, Revit, & Banerjee, 2004) at 10 dB SNR. Thus, there were 6 blocks in total (3 clear, 3 noise). Stimuli were presented binaurally at 75 dB_A_ SPL (noise at 65 dBA SPL). Further stimulus and task details as well as the behavioral data associated with this task are reported in (Bidelman, Mahmud, et al., 2019).

### 2.3 EEG recording

During the behavioral task, neuroelectric activity was originally recorded from 32 channels at standard 10-20 electrode locations on the scalp (Oostenveld & Praamstra, 2001). Preprocessing procedures are described fully in our previous reports (Bidelman et al., 2019; 2019b). Data were re-referenced off-line to a common average. Following ocular artifact correction (Picton et al., 2000) cleaned EEGs were then filtered (1-100 Hz; notched filter 60 Hz), epoched (-10-200 ms), and averaged in the time domain (described in Sect. 2.5) to derive ERPs for each stimulus condition per participant. Responses to non-targets (/ba/ and /pa/ tokens) were collapsed to reduce the dimensionality of the data. Infrequent /ta/ responses were not analyzed given their limited number of trials.

### 2.4 EEG source localization

We localized the sources of the scalp-EEG data by performing a distributed source analysis. We performed source localization in the MATLAB package Brainstorm (Tadel, Baillet, Mosher, Pantazis, & Leahy, 2011) using a realistic boundary element head model (BEM) volume conductor (M. Fuchs, Drenckhahn, Wischmann, & Wagner, 1998; Manfred Fuchs, Kastner, Wagner, Hawes, & Ebersole, 2002) standardized to the MNI brain (Mazziotta, Toga, Evans, Fox, & Lancaster, 1995). The BEM head model was implemented using the OpenMEEG module in Brainstorm (Gramfort, Papadopoulo, Olivi, & Clerc, 2010). A BEM is less prone to spatial errors than other existing head models (e.g., concentric spherical conductor) (Manfred Fuchs et al., 2002). Essentially, the BEM model parcellates the cortical surface into 15,000 vertices and assigns a dipole at each vertex with an orientation perpendicular to the cortical surface. From the pre-stimulus interval, the noise covariance matrix was estimated. We then used standard low-resolution brain electromagnetic tomography (sLORETA) to create inverse solutions (Pascual-Marqui, Esslen, Kochi, & Lehmann, 2002). We used Brainstorm’s default regularization parameters (regularized noise covariance = 0.1; SNR = 3.00). sLORETA provides a smoothness constraint that ensures the estimated current changes little between neighboring neural populations on the brain (Michel et al., 2004; Picton et al., 1999). Mean localization error for sLORETA is estimated to be 1.45 mm for 32 channels (Michel et al., 2004). From each single trail sLORETA brain volume, we extracted the time-course of source activity within regions of interest (ROI) defined by the Desikan-Killany (DK) atlas parcellation (Desikan et al., 2006). This atlas has 68 ROIs (e.g., LH: 34 ROIs, and RH: 34 ROIs). Subsequently, these time-courses were used as input to the SVM and stability selection to investigate *when* and *where* brain activity distinguishes NH and HI groups.

### 2.5 Feature extraction

Generally, averaging over more trials enhances the signal to noise ratio (SNR) of the ERPs by reducing EEG noise. Our dataset included ∼6000 trials per subject and condition (clear, noise) that can provide an adequate number of training and test examples using ERPs computed with different subsets of trials (without replacement). From each of the 68 ROIs of the DK atlas, we extracted source-level ERPs (i.e., mean activation within each ROI) averaged over randomly chosen 25, 50, 75, 100, and 125 trials without replacement. We then analyzed the ERP time courses using a sliding window basis (10 ms without overlap) across the whole epoch. Empirically, we found that responses averaged over 100 trials yielded the best classification results, providing a balance between classifier performance, computational speed, while also ensuring adequate SNR of the ERPs. 100 trial averages are therefore reported hereafter. The sliding window resulted in 21 (i.e., 210ms/10ms) ERP features (i.e., mean amplitude per window) for each ROI waveform, yielding 68*21=1428 features for each condition (e.g., clear and noise). These features were used as the input to the SVM classifier and stability selection coupled with SVM framework. As is common in classifiers, data were z-score normalized prior to classification and stability selection in order to ensure all features were on a common scale and range (Casale, Russo, Scebba, & Serrano, 2008).

### 2.6 SVM classification

Data driven multivariate analysis are a mainstay in modeling complex data and understand the relationship among all possible variables. Parameter optimized SVM classifiers are better candidate in building robust discriminative models with small sample sizes, which is common in human neuroimaging studies (Furey et al., 2000; Polat & Güneş, 2007). Classifier performance is greatly affected by the choice of kernel function, which can be used to map nonlinearly separable data to linearly separable space. Other tunable parameters (e.g., kernel, *C, γ*) also alter performance (Hsu, Chang, & Lin, 2003). As such, we used a grid search approach (range of *C* = 1e-2 to 1e2, and *γ* = 1e-2 to 7e-4) to find the optimal kernel, *C*, and *γ* values (kernels = ‘RBF’). We randomly split the data into training and test sets (80% and 20%, respectively) (Park, Luo, Parhi, & Netoff, 2011).

During the training phase, we fine-tuned the *C* and *γ* parameters to find optimal values for the classifier that maximally distinguished observations from the NH vs. HI group. The SVM learned the support vectors from the training data that comprised the attributes (e.g., ERPs amplitudes) and class labels (e.g., NH and HI). The resulting hyperplanes were fixed with maximum margin (e.g., maximum separation between the two classes) and used for predicting the unseen test data (by providing only the attributes but no class labels). Classification performance was calculated from standard formulas (accuracy, *F*1-score, and area under the curve (AUC)) (Saito & Rehmsmeier, 2015). AUC is a discriminating metric which describes the degree to which the model is capable of distinguishing between classes. An excellent model has AUC near to 1, meaning it has a good measure of separability. On the other hand, a poor model has AUC close to 0, meaning it has poor separability.

### 2.7 Stable feature selection (Stability selection)

A robust model should be complete enough to allow generalization and be easy to interpret. On the contrary, large numbers of feature variables (several thousand, as measured here) are susceptible to overfitting and can lead to models that are difficult to interpret. This requires selecting a set of the most salient discriminating features that are consistent across a range of model parameters. Feature selection is difficult, especially when the number of samples are small as compared to the number of features. Stability selection is a state-of-the-art feature selection method that works well in high dimensional or sparse problems based on Lasso (least absolute shrinkage and selection operator) (Meinshausen & Bühlmann, 2010; Yin, Li, & Zhang, 2017). Stability selection uses a Randomized Lasso algorithm, which works by resampling the training data set and computing feature scores on each resampling over the range of regularization parameters. Because stability selection includes an internal randomization technique (over many interactions), it yields a more reliable and consistent feature set than the conventional filtering and multivariate approaches. Stability selection can identify the most stable or relevant features from a large number of features, even if the necessary conditions required for the original Lasso method are violated (Meinshausen & Bühlmann, 2010).

In stability selection, a feature is considered to be more stable if it is more frequently selected over repeated subsampling of the data (Nogueira, Sechidis, & Brown, 2017). Basically, the Randomized Lasso randomly subsamples the training data and fits a L1-penalized logistic regression model to optimize the error. Lasso reduces the variance without substantially increasing bias during the subsampling process. Over many iterations, feature scores are (re)calculated. The features are shrunk to zero by multiplying the features’ co-efficient by zero while the stability score is lower. Surviving non-zero features are considered important variables for classification. Detailed interpretation and mathematical equations of stability selection are explained in Meinshausen & Bühlmann (2010). The stability selection solution is less affected by the choice of the initial regularization parameters. Consequently, it is extremely general and widely used in high dimensional data even when the noise level is unknown.

We considered sample fraction = 0.75, number of resampling = 1000, with tolerance = 0.01 (Al-Fahad, Yeasin, & Bidelman, 2019; Meinshausen & Bühlmann, 2010) in our implementation of stability selection. In the Lasso algorithm, the feature scores were scaled between 0 to 1, where 0 is the lowest score (i.e., irrelevant) and 1 is the highest score (i.e., most salient or stable feature). We estimated the regularization parameter from the data using the LARs algorithm (Efron, Hastie, Johnstone, & Tibshirani, 2004; Friedman, Hastie, & Tibshirani, 2010). Over 1000 iterations, Randomized Lasso provided the overall feature scores (0∼1) based on frequency a variable was selected. We ranked stability scores to identify the most important, consistent, stable, and invariant features (i.e., neural amplitudes across all ROIs and time) over a range of model parameters. We submitted these ranked features and corresponding class labels to an SVM classifier with different stability thresholds and observed the model performance. Based on the input stable features, SVM classified the group membership. The optimal stability threshold was selected corresponding to the maximum accuracy based on the AUC.

## 3. RESULTS

### 3.1 ERPs in HI vs. NH older adults

We first quantified the source (region-specific) ERPs of NH and HI listeners during clear and noise-degraded speech perception. For visualization purposes of the raw data, Figure 2 presents source waveforms for clear and noise-degraded speech within four selected ROIs reported in our previous study (Bidelman, Mahmud, et al., 2019). A detailed analysis of the source ERPs is reported elsewhere (Bidelman, Price, et al., 2019).

**Figure 2:**
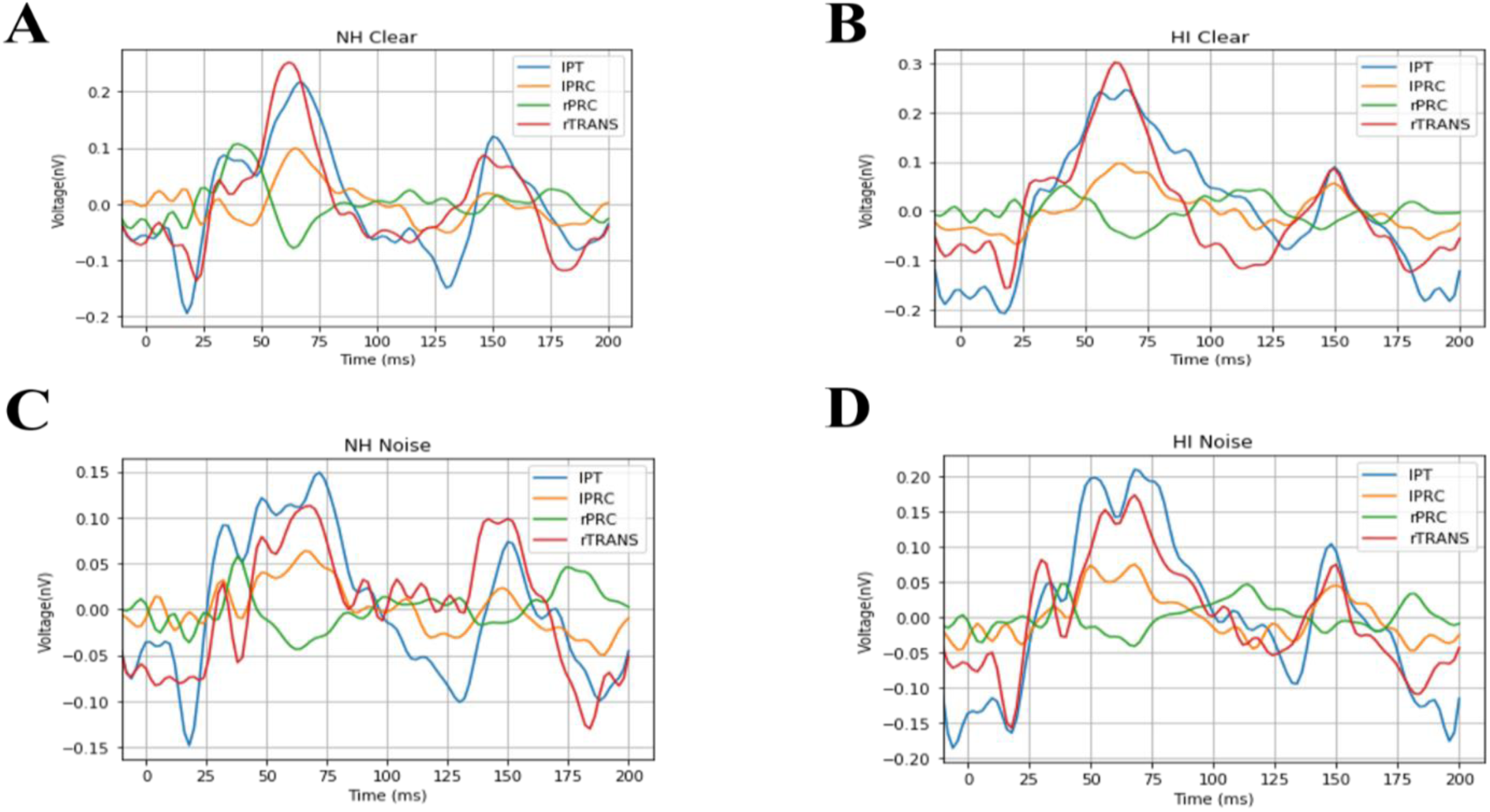
Source level ERPs for the NH and HI group in select ROIs. (A, B) Clear speech responses. (C, D) Noise-degraded speech responses. Note that traces are visualized here without baseline correction. This was accounted for in z-score normalization (i.e., mean centering) of all dependent measures prior to classifier analyses. NH, normal hearing; HI, hearing impaired; L, Left; R, Right; lPT, parstriangularis L; lPRC, precentral L; rPRC, precentral R; rTRANS, transverse temporal R.

### 3.2 SVM classification of hearing status using ERP features

We separately analyzed group classification performance using whole-brain source waveform data as well as each hemisphere (e.g., LH and RH) individually. We submitted ERP amplitudes and corresponding class labels to the SVM using a sliding window (10 ms) basis over the entire 210 ms epoch window (see Fig. 2). We used 5-fold cross-validation (Bhasin & Raghava, 2004) and carried out the grid search approach during the training period to determine the optimal parameters of the classifier. We then selected the best model and performance metrics from the predicted class labels, which were obtained from the unseen test data as well as true class labels. The time-varying SVM classifier accuracy is presented in Figure 3; maximum accuracy and its corresponding latency are shown in Figure 4 and summarized in Table 1.

**Table 1:** SVM classifier maximum performance (%) distinguishing hearing status (NH vs. HI)

| Speech stimulus | Measure | Whole brain features | LH features | RH features |
| --- | --- | --- | --- | --- |
| Clear | Accuracy | 82.03 | 80.83 | 78.64 |
|  | AUC | 81.18 | 80.16 | 78.13 |
|  | F1-score | 82.00 | 80.00 | 79.00 |
| Noise | Accuracy | 78.39 | 75.06 | 73.40 |
|  | AUC | 78.74 | 75.20 | 72.84 |
|  | F1-score | 79.00 | 75.00 | 74.00 |

**Figure 3:**
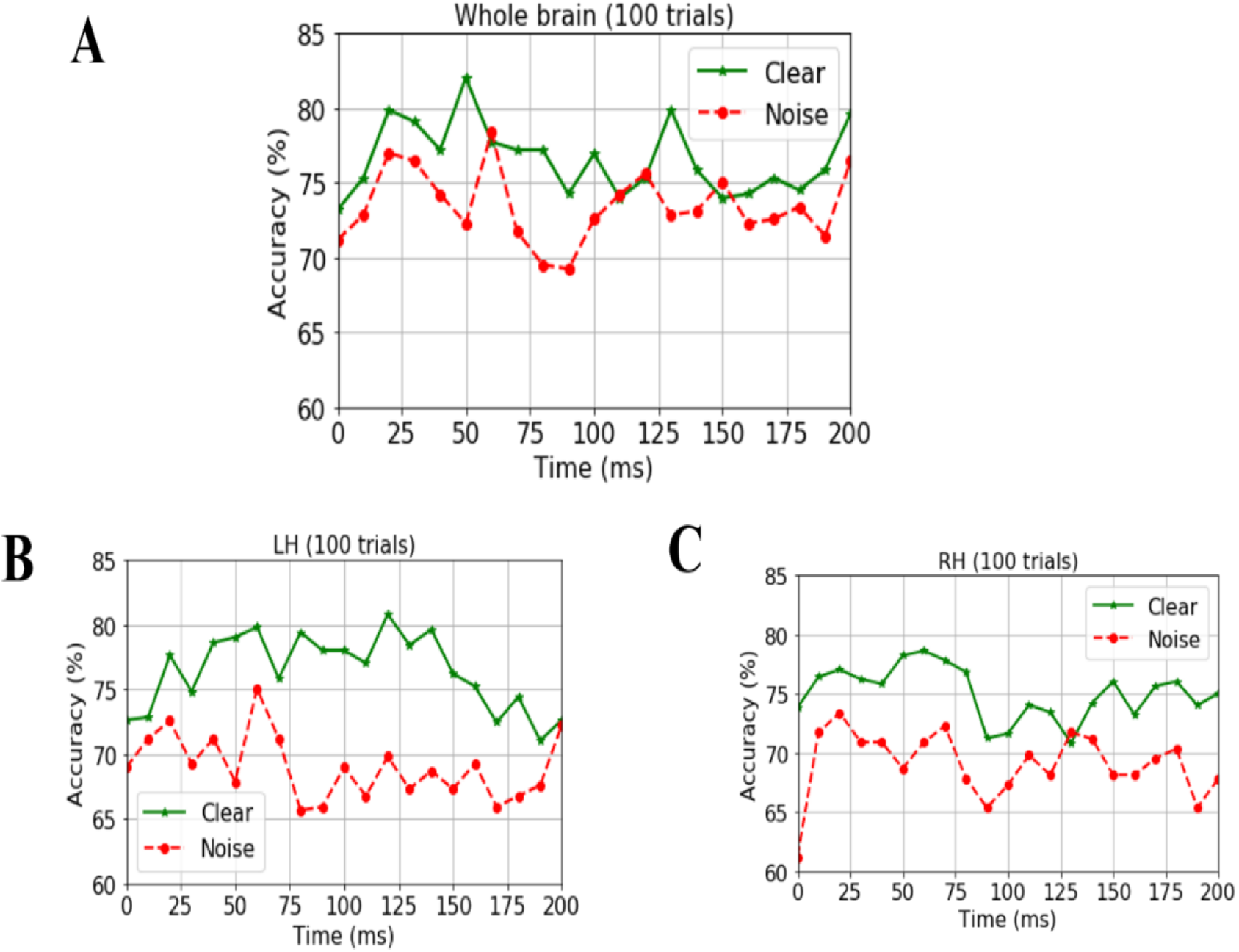
Time-varying classification of NH and HI groups with full brain vs. hemisphere-specific ERPs. Group classification accuracy from (A) whole-brain data (all 68 ROIs), (B) LH data alone, and (C) RH data alone. LH, left hemisphere; RH, right hemisphere.

**Figure 4:**
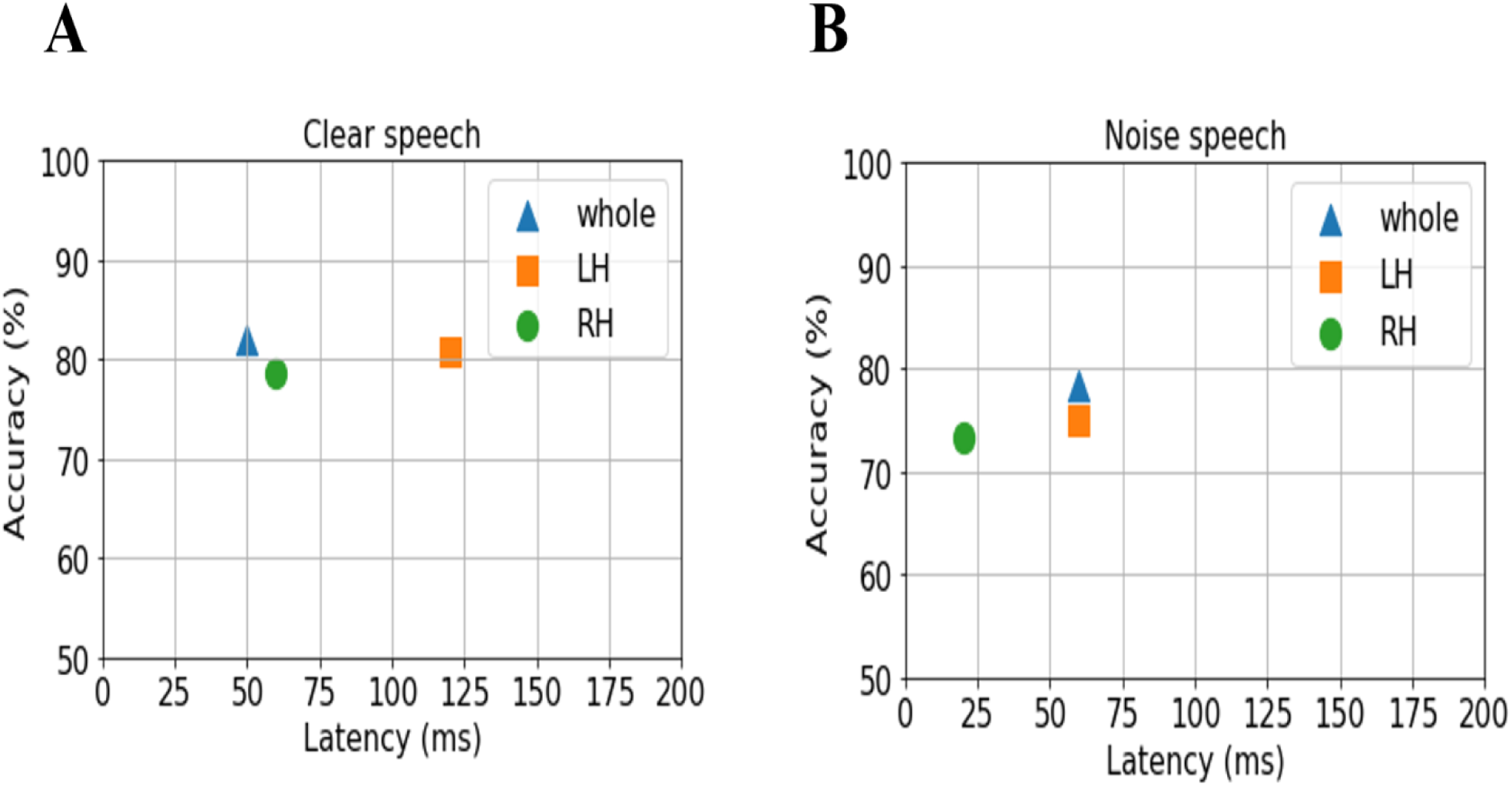
Maximum classifier accuracy and corresponding latency for distinguishing NH and HI using ERPs from the whole brain and LH vs. RH. (A) Clear speech responses. (B) Noise-degraded speech responses.

For ERP responses elicited by *clear* speech, the classifier conducted on full-brain data (all 68 ROIs) yielded a maximum accuracy in labeling groups of 82.03% at 50 ms. Classification was still quite accurate using LH (34 ROIs) responses alone (80.83%), but group segregation occurred at later latency at 120 ms. The poorest accuracy was obtained using RH (34 ROIs) features alone (78.64%) at a latency of 60 ms.

For ERP responses to *noise*-*degraded* speech, the maximum classifier accuracy was 78.39% at 60 ms using full-brain data. The LH features showed slightly lower accuracy (75.06%) than the whole brain at a latency of 60 ms. RH features provided the lowest accuracy (73.40% at 20 ms), among the three feature scenarios. Still, these group classification results are well above chance (i.e., 50%) and reveal the temporal dynamics of cortical speech activity robustly identifies older listeners with mild hearing loss.

### 3.3 Stability selection coupled with SVM

We used stability selection to identify the most important brain ROIs that can segregate groups with minimal computation and without overfitting. ERPs features were considered stable if they yielded higher stability scores at an 80% criterion level of classification performance (i.e., > 80% group separation). During pilot modeling, we roved stability thresholds which yielded different levels of classification performance. The effect of stability selection threshold on model performance is delineated in Figure 5A (clear) and Figure 5B (noise degraded). The histogram demonstrates the distribution of feature scores. The first line of x-axis represents the stability score (0 to 1); the second and third line represent the number and percentage of selected features under the corresponding bin; line four shows the number of cumulative unique brain ROIs up to the lower boundary of the bin. The semi bell-shaped solid black and red dotted lines of Figure 5 indicate the accuracy and AUC curve for different stability scores, respectively. In our stability selection analysis, the number of features represents ROI-specific source ERP amplitudes (in different time windows) and the number of unique ROIs represent functionally distinct brain areas of the DK atlas.

**Figure 5:**
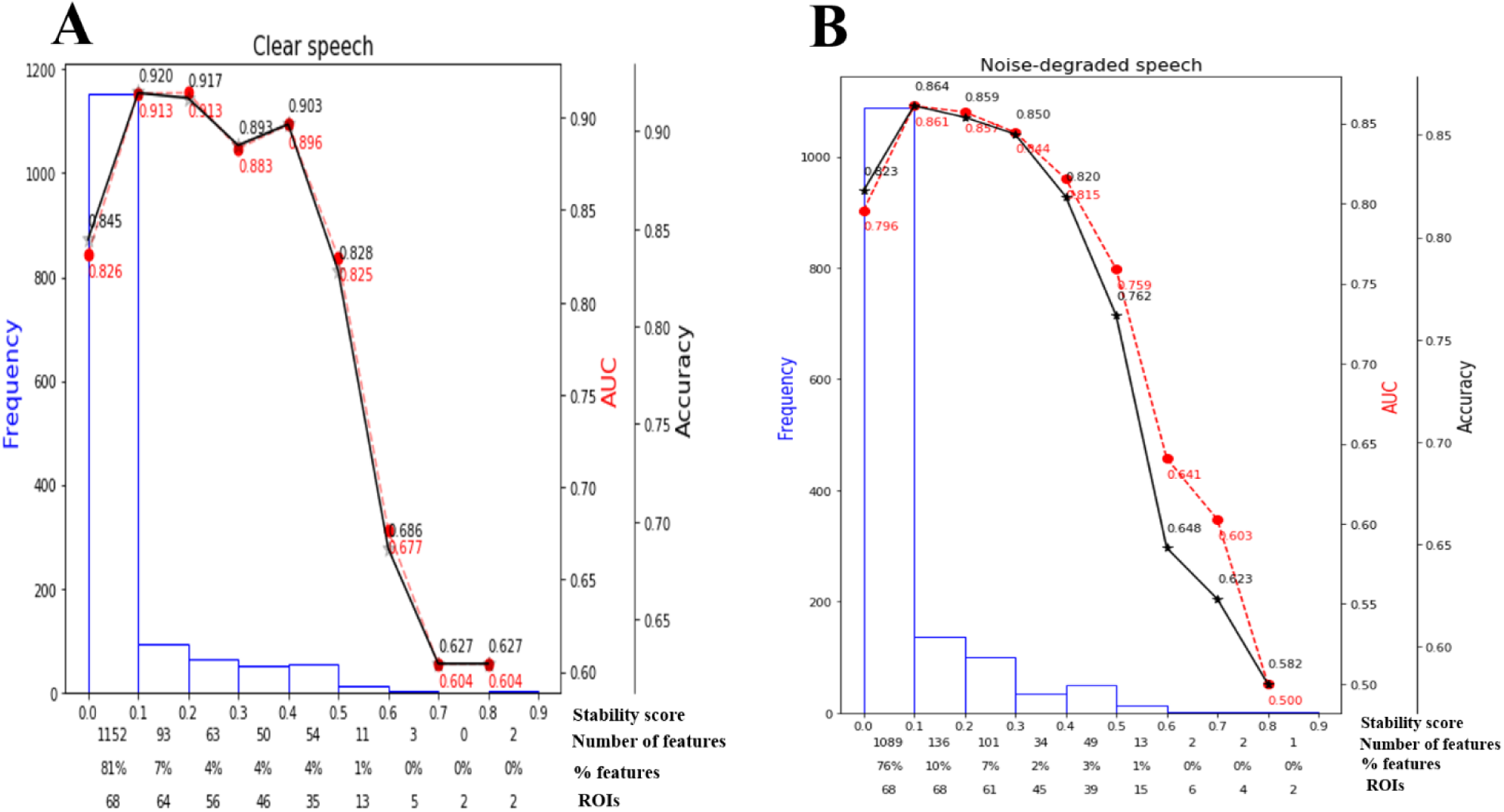
Effect of stability score threshold on model performance. The bottom of x-axis has four labels; *Stability score*, represents the stability score range of each bin (scores: 0∼1); *Number of features*, number of features under each bin; *% features*, corresponding percentage of selected features; *ROIs*, number of cumulative unique brain regions up to lower boundary of the bin. (A) Clear speech (B) noise-degraded speech.

The selected subset of features from the whole-brain identified via stability selection were then submitted to an SVM. For both clear and noise-degraded speech, the SVM classifier performance changed with the choice of stability threshold. We found that for clear speech detection, 81% of the features had scores (0 to 0.1) whereas 76% for noise-degraded speech detection. This means that the majority of ERP features were not selected among 1000 model iterations (i.e., carried near zero importance). Thus, 81% of the features were not related to segregating groups for clear speech, and 76% were irrelevant features for noise-degraded conditions.

Maximum accuracy in distinguishing groups was achieved using a stability score threshold of 0.10. Where the number of selected features 276 (19%) out of 1428 from 68 ROIs which yielded 92.0% accuracy (with AUC 91.3%, F1-score 92%) for clear speech responses; for noise degraded speech, stability selection selected 339 (24%) out of 1428 features from 68 ROIs, corresponding to 86.4% accuracy (with AUC 86.1%, F1-score 86%). Less than or greater than this optimal stability threshold the classifier showed poorer performance. Below the optimal threshold of 0.1, classifier performance was lower because irrelevant features were selected, whereas above the 0.1 threshold, relevant features to finishing hearing status were discarded.

Moreover, even when we selected a stability threshold of 0.7 (more conservative feature selection), clear speech responses could segregate groups with 62.3% accuracy using only two ROI. In contrast, noise-degraded speech yielded an accuracy of 62.7% with four ROIs. These results indicate that hearing status can be decoded still above chance levels using only a few brain regions engaged during speech perception. It is also notable that a larger number of brain ROIs were selected in noise-degraded speech perception as compared to clear speech perception corresponding to the same threshold and same accuracy.

Balancing these lax vs. strict models, we found that mid-level stability threshold of 0.5 segregated groups with an accuracy of 82.8% [AUC (82.5%) and F1-score (83.0)] by selecting only 16 features from 13 unique ROIs (reported in the Table 2) of the DK atlas. It was observed that accuracy degraded by 9% (from 92.0% to 82.8 %) from the optimal value but the number of features reduced dramatically from 276 to 16, stemming from only 13 (rather than 64) ROIs. For noise-degraded speech perception, only 19 features edges were selected from 15 unique ROIs and produced accuracy 76.2%, AUC 75.9% and F1-score 76.0% [i.e., accuracy degraded by 10 % from optimal accuracy (86.4%)] but well above chance level even in noise-degraded conditions. Thus, we considered a stability selection threshold of 0.5 which provided reasonable performance and less computation, but more critically, an interpretable network to describe neurophysiological speech processing. The network of brain ROIs (at 0.5 stability threshold) for clear and noise-degraded speech perception are shown in Figure 6–7 using the BrainO (Moinuddin, Yeasin, & Bidelman, 2019) visualization tool and detailed in Table 2.

**Table 2:**
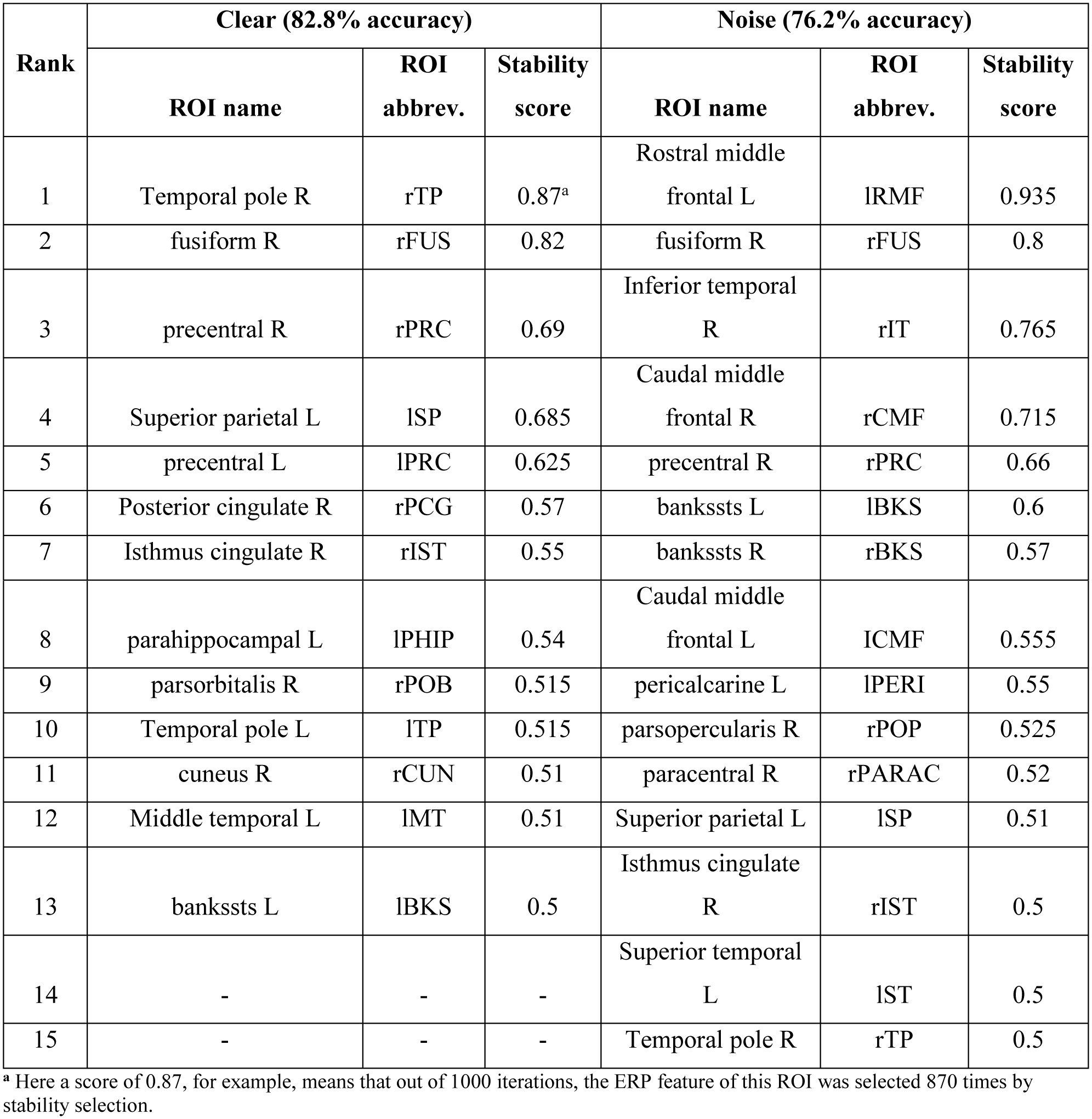
Most important brain regions (clear: 13 ROIs; noise: 15 ROIs; 0.50 stability threshold) distinguishing xsage-related hearing loss via EEG.

**Figure 6:**
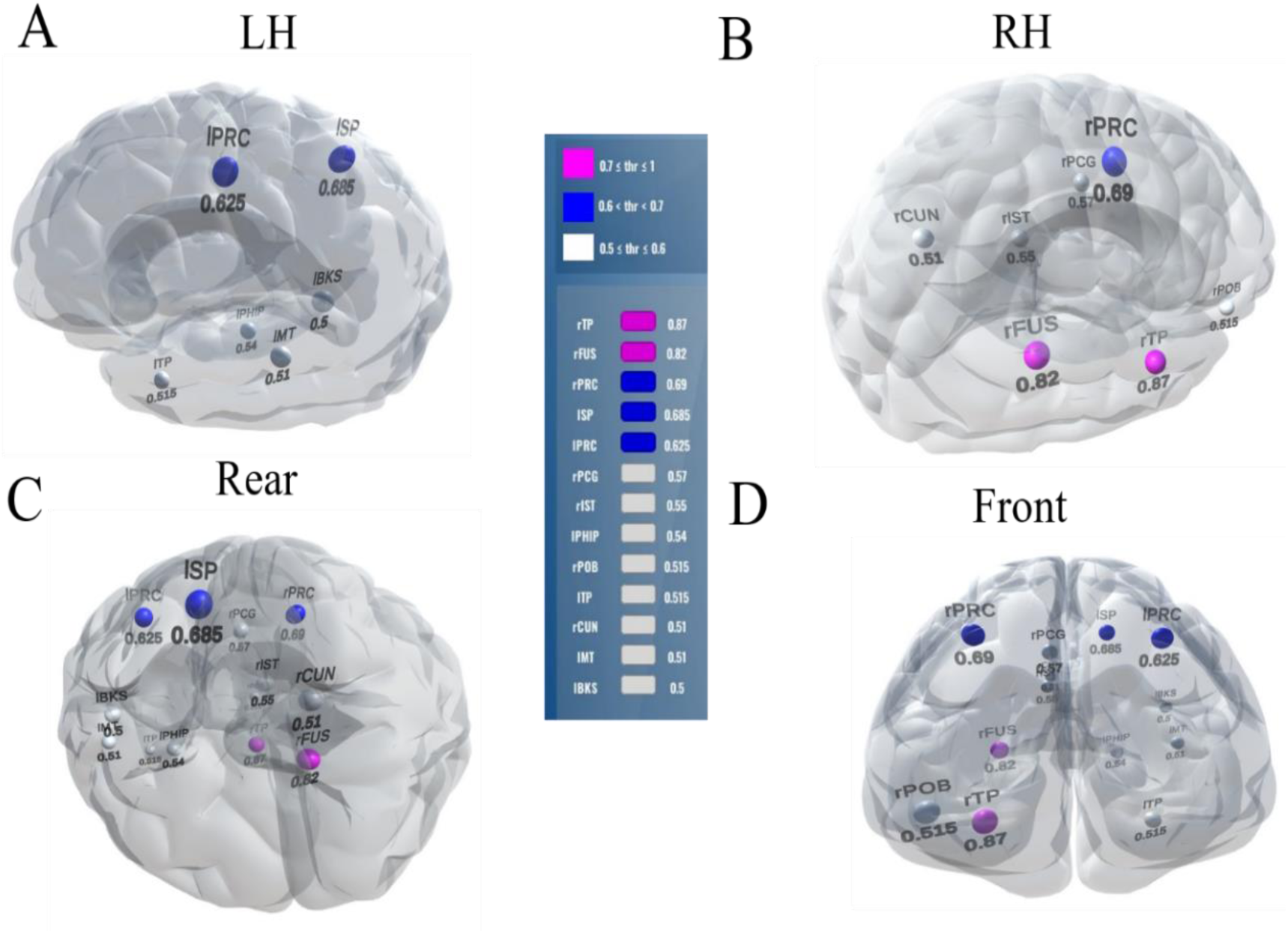
Stable (most consistent) neural network distinguishing NH and HI listeners during *clean* speech processing. Visualization of brain ROIs corresponding to 0.50 stability threshold (13 top selected ROIs which segregate groups at 82.8%) for clear speech perception. (A) LH (B) RH (C) Rear (D) Front. Stability score (color legend): (0.70 ≤ pink ≤1.0); (0.60 < blue < 0.70); (0.50 ≤ white ≤0.60).

**Figure 7:**
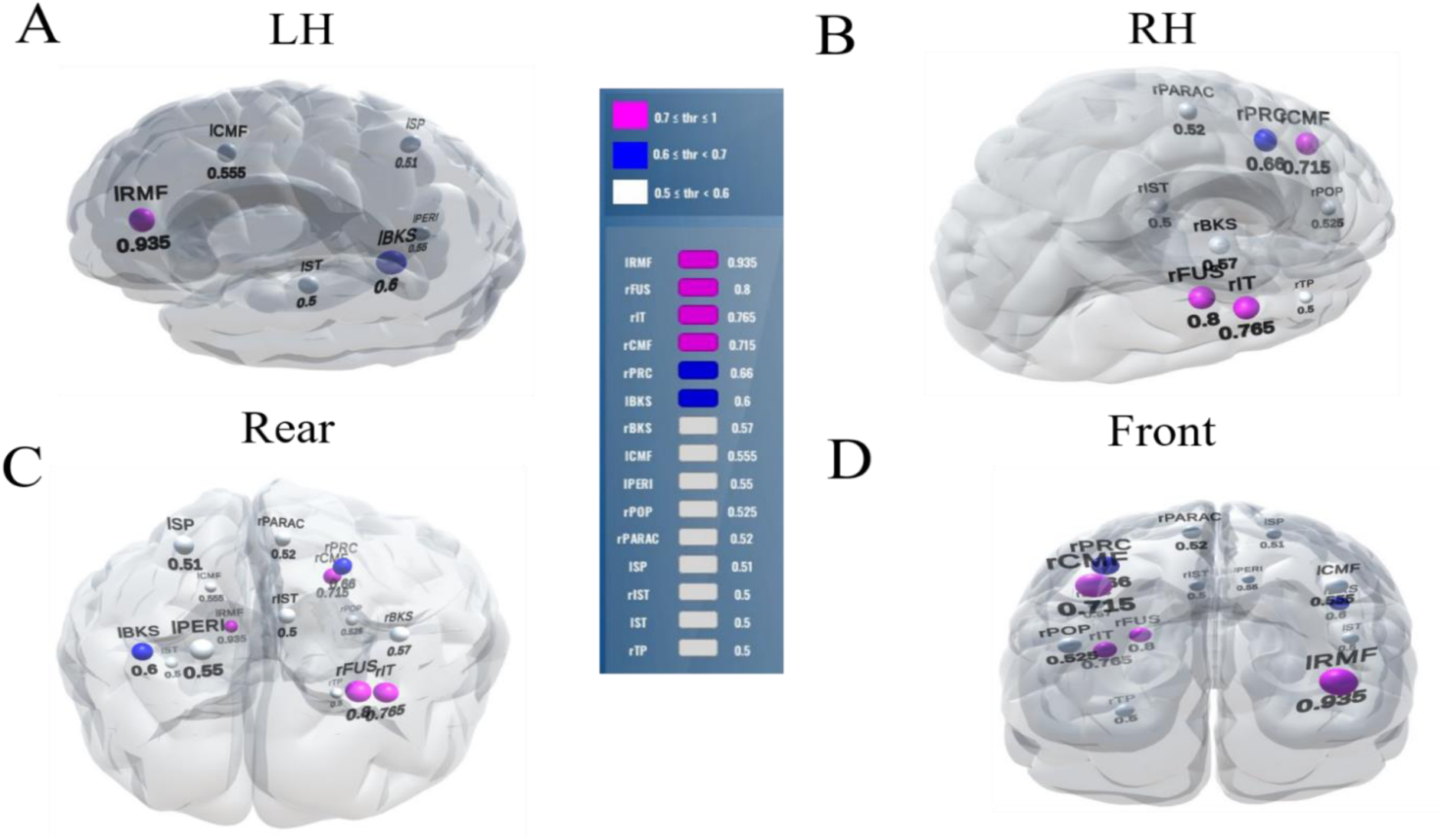
Stable (most consistent) neural network that distinguishes NH and HI listeners during *noise-degraded* speech processing. 15 top selected ROIs, 76% group classification. Otherwise as in Fig. 6.

## 4. DISCUSSION

In this study, we applied state-of-the-art machine learning techniques to EEG to decode the spatiotemporal dynamics of neural speech processing and identify *when* and *where* auditory brain activity is most sensitive to age-related declines in hearing.

### Hearing status is decoded early within the time-course of speech processing

Our data corroborate previous studies showing speech-ERPs are higher in amplitude for clear compared to noise-degraded speech detection and HI compared to NH listeners, consistent with the effects of hearing loss and background noise on auditory cortical responses (Alain, 2014; Bidelman, Price, et al., 2019; Bidelman et al., 2014). Extending previous work, we used these ERPs attributes in SVM classification to assess the time-course of the brain’s response to speech and their ability to segregate normal and hearing-impaired listeners. Among the three classification scenarios (e.g., whole-brain, LH, and RH analyses), whole-brain data provided the best group differentiation in the time frame of the P1 wave (∼50 ms). Additionally, LH activity provided better differentiation of groups’ speech processing than the RH.

Classification based on the sliding window analysis (full-brain level) showed maximum group decoding accuracy at 50 ms for clear speech, whereas noise-degraded speech, maximum accuracy was observed later at 60 ms. These results suggest that the P1 can be a useful attribute to segregate NH and HI listeners’ neurophysiological processing of speech but also depends on the clarity (i.e., SNR) of the speech signal. Since the P1 wave is thought to be generated by the thalamus and primary auditory cortex (Eggermont, Ponton, Don, Waring, & Kwong, 1997; Erwin & Buchwald, 1987; Jang et al., 2010; Liegeois-Chauvel, Musolino, Badier, Marquis, & Chauvel, 1994; McGee & Kraus, 1996), this suggests early hearing loss in older adults changes auditory processing in early sensory regions. Furthermore, for noise-degraded speech, maximum group segregation was delayed 10 ms relative to clear speech. The later decoding for degraded speech is perhaps expected due to inherent masking of the signal, which weakens the neural representation for speech, decreases the amplitude, and prolongs the latency of the ERPs. Previous studies have indeed shown that neural responses are significantly influenced by noise in the midbrain (Anderson, Skoe, Chandrasekaran, & Kraus, 2010; Burkard & Sims, 2002; Ding & Simon, 2013; Presacco et al., 2016) and cortex (Billings, McMillan, Penman, & Gille, 2013; Billings, Penman, McMillan, & Ellis, 2015). Thus, the delay in maximum decoding accuracy we find in the SIN condition is consistent with prior work. Moreover, the better performance by LH compared to RH activity in distinguishing groups is consistent with the dominance of LH in auditory language processing (Frost et al., 1999; Tervaniemi & Hugdahl, 2003; Zatorre, Evans, Meyer, & Gjedde, 1992).

### Degraded speech processing requires more (right hemisphere/frontal) neural resources than clear speech processing

We extend previous neuroimaging studies by demonstrating the most stable, consistent, and invariant functional brain regions supporting age-related speech and SIN processing using a data-driven approach (stability selection coupled with SVM). Stability selection with randomized Lasso on full-brain neural data identified the most important brain regions associated with hearing loss over a range of model parameters. Our analysis revealed the most stable brain regions could segregate groups at >80% accuracy from the ERP features alone corresponding to an optimal stability score (0.1). Group segregation was still reasonably accurate using a more stringent stability score of 0.5 which identified a sparser subset of brain regions that described age-related declines in speech processing (Table 2).

For clear speech perception, stability selection identified 13 regions, including five ROIs from temporal lobe (temporal pole R, fusiform R, temporal pole L, parahippocampal L, temporal pole L) and left auditory cortex (bankssts L), three regions from frontal lobe (precentral R, precentral L, parsorbitalis R), three from parietal lobe (superiorparietal L, posteriorcingulate R, isthmuscingulate R), one region from occipital lobe (cuneus R). For noise-degraded speech perception, four important regions emerged in the temporal lobe (fusiform R, inferior temporal R, superior temporal L, temporal pole R) including superior temporal sulcus (bankssts L, bankssts R), six regions from frontal lobe (rostral middle frontal L, caudal middle frontal R, precentral R, caudal middle frontal L, parsopercularis R, paracentral R), two from partial lobe (superiorparietal L, isthmuscingulate R), and one region from occipital lobe (pericalcarine L).

Among the two networks identified via stability selection, six regions were common for clear and noise-degraded speech perception. One of these, the right temporal pole is part of the auditory ventral pathway which plays a role in the coding, representation, and perception of nonspatial attributes of sound including auditory identity (Poremba et al., 2004; Tsunada, Lee, & Cohen, 2011). Recruitment of precentral gyrus (precentral R, precentral L) in our speech tasks is probably also anticipated given the role of primary motor areas in phoneme processing, particularly in noise (Du et al., 2014; Hickok, 2014). Superior temporal areas—especially in left hemisphere (l/rBKS, lSTS)—were also recruited, consistent with their role in the phonological network supporting speech perception and processing (Hickok & Poeppel, 2007).

Interestingly, for noise-degraded speech perception, we found several non-overlapping regions including dorsolateral prefrontal cortex (rostral middle frontal, caudal middle frontal). The additional recruitment of these areas in noise most probably reflects compensatory processing associated with working memory and higher-order speech processing (Gabrieli, Poldrack, & Desmond, 1998; Wong, Parsons, Martinez, & Diehl, 2004; Zatorre et al., 1992) that would necessarily need to be engaged during the more complex listening demands of noise. Perhaps involvement of the other non-overlapping regions also aids noise-degraded speech perception, in a yet unknown way. For example, we speculate that the dual involvement of the pericalcarine and superiorparietal areas in the noise condition may reflect a form of visual or motor imagery listeners use as a strategy to cope with task difficulty (Ganis, Thompson, & Kosslyn, 2004; Tian & Poeppel, 2012). It is noticeable that for clear speech, about half (6/13 = 46%) of the stable regions were from LH. However, LH involvement was reduced for noisy speech perception (6/15 = 40%) which was paralleled by stronger RH involvement (i.e., 9/15 = 60% stable regions were right lateralized). These findings are broadly consistent with previous neuroimaging studies demonstrating that noise-degraded speech perception requires additional RH brain regions to compensate for the impoverished acoustic signal (Bidelman & Howell, 2016; Mudar & Husain, 2016; Shtyrov et al., 1998; Shtyrov, Kujala, Ilmoniemi, & Näätänen, 1999).

Collectively, our findings show that frontal brain regions are associated with noise-degraded speech perception. Previous neuroimaging studies (Du et al., 2016; Erb & Obleser, 2013; Peelle et al., 2009; Vaden et al., 2015; Wong et al., 2009) have similarly demonstrated that noise-degraded speech perception increases recruitment of frontal brain regions (i.e., upregulation of frontal areas). Our data driven approach corroborates previous studies by confirming more brain regions are allocated to process acoustically-degraded compared to clear speech.

## 5. CONCLUSION

We developed a robust and efficient computational framework to investigate *when* and *where* cortical brain activity segregates NH and HI listeners by using a data-driven approach. Data driven machine learning analysis showed that the P1 wave of the auditory ERPs (∼50 ms) robustly distinguished NH and HI groups, revealing speech-evoked neural responses are highly sensitive to age-related hearing loss. Our results further suggest that identifying listeners with mild hearing impairment based on their EEGs is also more robust when using LH compared to RH features of brain activity, particularly under listening conditions that tax the auditory system (i.e., noise interference). From stability selection and SVM classifier analyses, we identified sparse (<15 regions) yet highly robust networks that describe older adults’ speech processing. Yet, we found 3-5% more neural resources are required to distinguish hearing-related declines in speech processing in noise, particularly those in the frontal lobe and right hemisphere.

## Acknowledgements

Requests for data and materials should be directed to GMB. This work was partially supported by the National Institutes of Health (NIH/NIDCD R01DC016267) and Department of Electrical and Computer Engineering at U of M.

